# Comparative genomics of beetle-vectored fungal pathogens reveals a reduction in genome size and independent evolution of pathogenicity of two tree pathogens

**DOI:** 10.1101/093856

**Authors:** Taruna A Schuelke, Anthony Westbrook, Keith Woeste, David C. Plachetzki, Kirk Broders, Matthew D. MacManes

## Abstract

- *Geosmithia morbida* is an emerging fungal pathogen which serves as a paradigm for examining the evolutionary processes behind pathogenicity because it is one of two known pathogens within a genus of mostly saprophytic, beetle-associated, fungi. This pathogen causes thousand cankers disease in black walnut trees and is vectored into the host via the walnut twig beetle. *G. morbida* was first detected in western US and currently threatens the timber industry concentrated in eastern US.
- We sequenced the genomes of *G. morbida* and two non-pathogenic *Geosmithia* species and compared these species to other fungal pathogens and nonpathogens to identify genes under positive selection in *G. morbida* that may be associated with pathogenicity.
- *G. morbida* possesses one of the smallest genomes among the fungal species observed in this study, and one of the smallest fungal pathogen genomes to date. The enzymatic profile is this pathogen is very similar to its relatives.
- Our findings indicate that genome reduction is an important adaptation during the evolution of a specialized lifestyle in fungal species that occupy a specific niche, such as beetle vectored tree pathogens. We also present potential genes under selection in *G. morbida* that could be important for adaptation to a pathogenic lifestyle.

## Introduction

Uncovering the specific genetic and molecular events behind the evolution of novel traits such as pathogenicity in fungal species has been a long-standing objective of pathologists. *Geosmithia* (Ascomycota: Hypocreales), a genus first proposed in 1979 for fungi that were formerly placed in genus *Penicillium* (Pitt, 1979), serves a paradigm for examining the processes contributing to the evolution of pathogenicity. *Geosmithia* species are filamentous fungi that most commonly associate with phloeophagous bark beetles (Kolarik *et al.*, 2005; Kolarik *et al.*, 2011), although some *Geosmithia* fungi, such as *G. eupagioceri* and *G. microcorthyli*, are known to affiliate with ambrosia beetles (Kolarik & Jankowiak 2013). *Geosmithia* species and their beetle associates occupy a variety of hosts, including pines, oaks, junipers, *and* walnut trees (Kolarik *et al.*, 2007; Kolarik & Kirkendall 2010; Kolarik & Jankowiak 2013). The ecology and diversity of symbiosis between these fungi and their beetle associates is poorly understood, but investigators are beginning to explore such relationships (Kolarik *et al.*, 2007; Kolarik & Jankowiak 2013). While most species in *Geosmithia* are saprotrophic, two species were recently determined to be pathogenic—*G. pallida* (Lynch *et al.*, 2014) and *G. morbida* (Tisserat *et al.*, 2009), on coast live oak *(Quercus agrifolia)* and black walnut *(Juglans nigra)*, respectively. However, both of these species live saprophytically in association with bark beetles and other tree hosts. It is still unclear what mechanisms allow these species of *Geosmithia* to be pathogenic to a new host while other members of the genus remain saprobes.

*Geosmithia morbida* causes thousand cankers disease (TCD) in *Juglans nigra* (eastern black walnut). Although no evidence of TCD has been detected in other *Juglans* to date, several species, such as *J. californica, J. cinerea, J. hindsii, J. regia*, are also susceptible to the pathogen (Utley *et al.*, 2013). The fungus is most often vectored into its hosts by *Pityophthorus juglandis*, commonly known as the walnut twig beetle (WTB) (Kolarik *et al.*, 2011). Unusual mortality of *J. nigra* was first noted in Colorado, US in 2001. Since then, nine western states (CO, WA, OR, ID, NV, UT, CA, NM, AZ) and seven eastern states (PA, OH, IN, MD, VA, TN, NC) have reported TCD in one or more locations (Zerillo *et al.*, 2014). This increase in TCD is likely a consequence of the expansion of WTB’s geographic range. WTB was present in only four counties of California, Arizona and New Mexico in the 1960s, however, as of 2014, the beetle has been detected in over 115 counties in the western and eastern US (Rugman-Jones *et al.*, 2014).

The origin of this pathogen is not clear. However, it has been hypothesized that *G. morbida* may have undergone a host shift from *J. major* (Arizona black walnut) to a more naïve host, *J. nigra*, because the fungus does not cause disease in the Arizona black walnut, and neither WTB nor *G. morbida* were observed in the native range of *J. nigra* until 2010 (Zerillo *et al.*, 2014). *J. nigra* is not indigenous to western US but was planted throughout the region as an ornamental species. An alternative prediction based on *G. morbida* population genetic data is that the origin of *G. morbida* and WTB are the walnut populations of southern California, where the pathogen has been isolated from both healthy and diseased *J. californica* trees (Zerillo *et al.*, 2014).

Early symptoms of infection by *G. morbida* include yellowing, wilting and thinning of the foliage followed by branch dieback and tree death within 2–3 years after the initial infestation (Tisserat *et al.*, 2009; Kolarik *et al.*, 2011). Little is known about the specific means *G. morbida* employs for initiating and maintaining the infection, or what benefits, if any, the fungus imparts to the WTB vector. However, previous studies have demonstrated that fungal pathogens that occupy ecological niches similar to *G. morbida* must be capable of enduring and combating toxic host environments used by plants to resist infection. For instance, *Grosmannia clavigera*, a fungal symbiont of the mountain pine beetle *(Dendroctonus ponderosae)*, can detoxify metabolites such as terpenoids and phenolics produced by the host as defense mechanisms (DiGuistini *et al.*, 2011).

We recently developed a reference genome of *Geosmithia morbida* that consisted of 73 scaffolds totaling 26.5 Mbp in length (Schuelke *et al.*, 2016). This genome represents one of the smaller fungal tree pathogen genomes reported to date. Rapid changes in genome size have accompanied dramatic biological changes in newly emerged fungal and oomycete species (Raffaele & Kamoun 2012; Adhikari *et al.*, 2013). In fungi, a link has been observed between genome expansion and evolution of pathogenicity (Raffaele & Kamoun 2012). Genome expansions were associated with parasitism in general and increased pathogenicity and virulence in several fungal lineages (Spanu *et al.*, 2010). Previous genome sequencing of *Geosmithia morbida* (Schuelke *et al.*, 2016) showed that this newly emerged fungal pathogen has a smaller genome than several of its closely related nonpathogenic relatives in the Hypocreales. Hence, *Geosmithia* has taken an evolutionary path to pathogenicity that has not been characterized previously in plant-associated fungi.

The evolution of pathogenicity via genome size reduction is not understood, and although it is contrary to our current expectation of how pathogenicity develops in non-pathogens, it might be common, particularly in beetle-associated symbionts. In fact, in the last decade several beetle-associated fungi have emerged as plant pathogens, including *Grosmannia clavigera*, which is also a tree infecting, beetle-vectored fungus, and its genome size is only 29.8 Mb (Diguistini *et al.*, 2011). The arrival of new pathogens, frequently referred to as Black Swan events due to their perceived unpredictability, represent a significant threat to native and agriculturally important tree species (Pleotz *et al.*, 2013). Thus, beetle-associated symbionts that have switched to pathogens represent excellent models for investigating the evolution of pathogenicity and its relationship to genome size. Although the genus *Geosmithia* is distributed worldwide, *G. morbida*, and more recently, *G. pallida*, are the first members of the genus to be described as plant pathogens among the 60 known nonpathogenic species (Kolarik & Kirkendall 2010; Kolarik *et al.*, 2011; Lynch *et al.*, 2014).

In this work, we compare the reference genome of the pathogenic and host specific species *G. morbida* with two closely related non-pathogenic generalist species, *G. flava* and *G. putterillii.* Based on this comparison, we identify genes under positive selection that may be involved in the specialization of a pathogenic life strategy that depends on a single beetle vector and a narrow, but potentially expanding, host range. We also present a species phylogeny estimated using single-copy orthologs that confirms the placement of *Geosmithia* species in the order Hypocreales, and that their closest fungal relative is *Acremonium chrysogenum.* The primary goal of this study was to gain insight into the evolution of pathogenicity within *G. morbida.* We also investigated the presence of convergent evolution in *G. morbida* and *Grosmannia clavigera*, two tree pathogens vectored into their hosts via beetle symbionts.

## Materials and Methods

### DNA extraction and sequencing

The CTAB method delineated by the Joint Genome Institute was used to extract DNA for genome sequencing from lyophilized mycelium of *Geosmithia flava* and *Geosmithia putterillii* (Kohler *et al.*, 2011). Table 1 lists genetic, geographic, and host information for each *Geosmithia* species used in this study. Total DNA concentration was measured with Nanodrop, and DNA sequencing was conducted at Purdue University Genomics Core Facility in West Lafayette, Indiana. DNA libraries were prepared using the paired-end Illumina Truseq protocol and sequenced on an Illumina HiSeq 2500 using a single lane. Mean insert sizes for *G. flava* and *G. putterillii* were 477bp and 513bp, correspondingly. The remaining sequencing statistics can be found in Table 2.

**Table 1.**
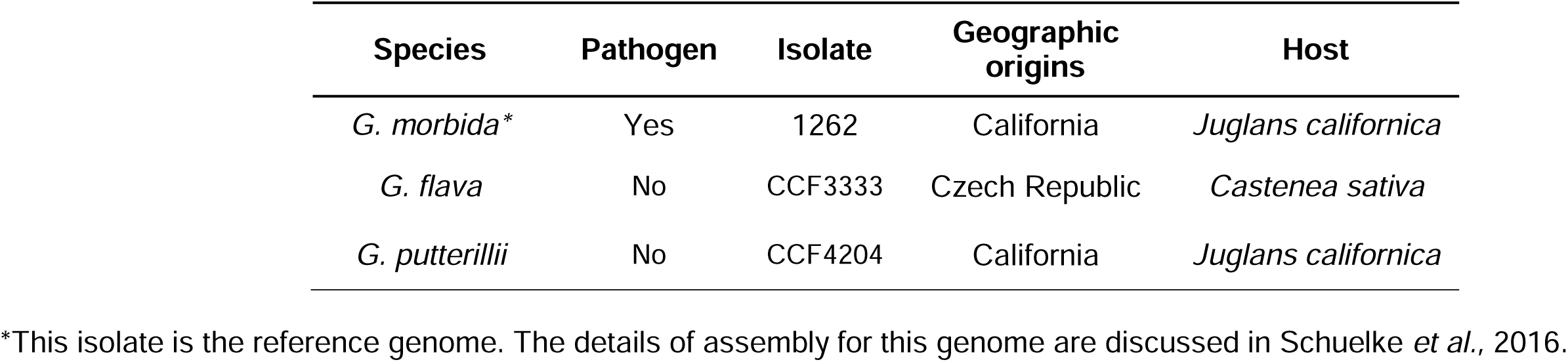
Species, geographic origins, and host information for *Geosmithia morbida*, *Geosmithia flava*, and *Geosmithia putterillii*.

| Species | Pathogen | Isolate | Geographic origins | Host |
| --- | --- | --- | --- | --- |
| *G. morbida* * | Yes | 1262 | California | *Juglans californica* |
| *G. flava* | No | CCF3333 | Czech Republic | *Castanea sativa* |
| *G. putterillii* | No | CCF4204 | California | *Juglans californica* |

\*This isolate is the reference genome. The details of assembly for this genome are discussed in Schuelke *et al.*, 2016.

**Table 2.** Statistics for sequence data from isolates of *Geosmithia morbida*, *Geosmithia flava* and *Geosmithia putterillii*.

| Species | Total read pairs |  | Est. coverage |  |
| --- | --- | --- | --- | --- |
| *G. morbida* | 14,013,863* | 20,674,289** | 109x* | 160x** |
| *G. flava* | 16,183,281 |  | 102x |  |
| *G. putterillii* | 19,711,745 |  | 131x |  |

\*These values are for paired-end read data for *G. morbida* from Schuelke *et al.*, 2016. \*\*These values are for mate-pair read data for *G. morbida* from Schuelke *et al.*, 2016.

### Preprocessing sequence data

The raw paired-end reads for *G. flava* and *G. putterillii* were corrected using BFC (version r181) (Li 2015). BFC utilizes a combination of hash table and bloom-filter to count *k*-mers for a given read and correct errors in that read based on the *k*-mer support. Because BFC requires interleaved reads as input, khmer 1.1 was leveraged to interleave as well as split the paired-end reads before and after the error correction stage, respectively (Crusoe *et al.*, 2015). Next, low quality bases and adapters in error corrected reads were trimmed with Trimmomatic, version 0.32, using a Phred threshold of 4 (Bolger *et al.*, 2014).

### Assembly construction

Genome assemblies were constructed with ABySS 1.9.0 using four k-mer sizes (61, 71, 81, and 91) (Simpson *et al.*, 2009). The resulting assemblies were evaluated using BUSCO (v1.1b1) (Simão *et al.*, 2015), which assess completeness based on the presence of universal single-copy orthologs within fungi. Length-based statistics were generated with QUAST v2.3 (Gurevich *et al.*, 2013). Final assemblies were manually chosen based on length-based and genome completeness statistics. Furthermore, the raw reads of *G. flava* and *G. putterillii* were mapped back to their corresponding genomes using BWA version 0.7.9a-r786 (Li & Durbin, 2009) to assess the quality of the chosen assemblies.

### Structural and Functional Annotation

We utilized the automated annotation software Maker version 2.31.8 (Cantarel *et al.*, 2008) to functionally annotate the genomes of *G. flava* and *G. putterillii.* We used two of the three gene prediction tools available within the pipeline SNAP (released 2013, Korf 2004) and Augustus 2.5.5 (Stanke *et al.*, 2006). SNAP was trained using gff files generated by CEGMA v2.5 (a program similar to BUSCO) (Parra *et al.*, 2007). Augustus was trained with *Fusarium solani* protein models (v2.0.26) downloaded from Ensembl Fungi (Kersey *et al.*, 2016). The protein sequences generated by the structural annotation were blasted against the Swiss-Prot database (Boutet *et al.*, 2016) to functionally annotate the genomes of *G. flava* and *G. putterillii*.

### Assessing repetitive elements profile

To evaluate the repetitive elements profile of *G. flava* and *G. putterillii*, we masked the interspersed repeats within the assembled genomes with RepeatMasker 4.0.5 (Smit *et al.*, 1996) using the sensitive mode and default values as arguments.

### Identifying putative genes involved in host-pathogen interactions

To search for putative genes contributing to pathogenicity, we conducted a BLASTp (v2.2.28+) (Altschul *et al.*, 1990) search with an e-value threshold of 1e-6 against the PHI-base 4.0 database (Winnenburg *et al.*, 2006) that includes known genes implicated in pathogenicity. Additionally, we identified proteins that contain signal peptides and lack transmembrane domains in each *Geosmithia* species as well as their close relative *Acremonium chrysogenum* with SignalP 4.1 and TMHMM 2.0 using default parameters (Krogh *et al.*, 2001; Peterson *et al.*, 2011).

### Identifying species specific genes

To identify unique genes present in *Geosmithia* morbida, we performed an all-versus-all BLASTp search among the three *Geosmithia* species and *A. chrysogenum* with Orthofinder version 0.3.0 (Emms & Kelly 2015). Using a <u>custom Python script</u>, we analyzed homology among the four fungal species.

### Identifying carbohydrate-active proteins and peptidases

To identify enzymes capable of degrading carbohydrate molecules in species belonging to Hypocreales and *G. clavigera*, we performed a HMMER 3.1b1 (Eddy 1998) search against the CAZy database (Lombard *et al.*, 2014) released July 2015 and filtered the results following the developer’s recommendations. Lastly, we profiled the proteolytic enzymes present in species using the *MEROPS* database 10.0 (Rawlings *et al.*, 2016).

### Phylogenetic analysis

#### Taxon Sampling

In order to determine phylogenetic position of *Geosmithia*, we combined the predicted peptide sequences from three *Geosmithia* species described here with the predicted peptide sequences of an additional 17 fungal genomes that represent the breadth of pathogens and non-pathogens within Ascomycota. Our dataset contained eleven pathogens and nine non-pathogens (Table 3).

**Table 3.** Fungal species used for phylogenetic analysis in this study. The species in bold were utilized for positive selection analysis.

| Species | Class | Order | Ecological role | Download source | References |
| --- | --- | --- | --- | --- | --- |
| ***Geosmithia morbida*** | Sordariomycetes | Hypocreales | Pathogen | - | Schuelke *et al.*, 2016 |
| ***Geosmithia flava*** | Sordariomycetes | Hypocreales | Non-pathogen | - | - |
| ***Geosmithia putterillii*** | Sordariomycetes | Hypocreales | Non-pathogen | - | - |
| ***Acremonium chrysogenum*** | Sordariomycetes | Hypocreales | Beneficial | FungalEnsembl | Terfehr *et al.*, 2014 |
| ***Stanjemonium griseum*** | Sordariomycetes | Hypocreales | Saprotrophic | JGI | Used with permission |
| *Trichoderma virens* | Sordariomycetes | Hypocreales | Mycoparasite | JGI | Kubicek *et al.*, 2011 |
| ***Trichoderma reesei*** | Sordariomycetes | Hypocreales | Saprotrophic | FungalEnsembl | Martinez *et al.*, 2008 |
| *Escovopsis weberi* | Sordariomycetes | Hypocreales | Mycoparasite | EnsemblGenomes | de Man *et al.*, 2016 |
| *Ustilagoidea virens* | Sordariomycetes | Hypocreales | Biotrophic pathogen | FungalEnsembl | Zhang *et al.*, 2014 |
| *Cordyceps militaris* | Sordariomycetes | Hypocreales | Insect pathogen | FungalEnsembl | Zheng *et al.*, 2011 |
| ***Myrothecium inundatum*** | Sordariomycetes | Hypocreales | Saprotrophic | JGI | Used with permission |
| *Fusarium solani* | Sordariomycetes | Hypocreales | Necrotrophic pathogen | FungalEnsembl | Coleman *et al.*, 2009 |
| *Fusarium graminearum* | Sordariomycetes | Hypocreales | Necrotrophic pathogen | FungalEnsembl | Trail *et al.*, 2003; Cuomo *et al.*, 2007; Ma *et al.*, 2010 |
| *Ceratocystis platani* | Sordariomycetes | Microascales | Pathogen | FungalEnsembl | Belbahri 2015 |
| ***Neurospora crassa*** | Sordariomycetes | Sordariales | Saprotrophic | FungalEnsembl | Galagan *et al.*, 2003 |
| ***Chaetomium globosum*** | Sordariomycetes | Sordariales | Saprotrophic | JGI | Berka *et al.*, 2011 |
| ***Grosmannia clavigera*** | Sordariomycetes | Ophiostomatales | Pathogen | FungalEnsembl | DiGuistini *et al.*, 2011 |
| *Eutypa lata* | Sordariomycetes | Xylariales | Pathogen | JGI | Blanco-Ulate *et al.*, 2013 |
| *Botrytis cinerea* | Leotiomycetes | Helotiales | Necrotrophic pathogen | FungalEmsebl | Amselem *et al.*, 2011,<br>Staats & van Kan 2012 |
| ***Oidiodendron maius*** | Leotiomycetes | Incertae sedis | Mycorrhizal | JGI | Kohler *et al.*, 2015 |

#### Inferring Orthology

Orthologous peptide sequences among the 20 fungal genomes were determined using OrthoFinder version 0.3.0 (Emms & Kelly 2015). OrthoFinder performs an all-versus-all BLASTp (2.2.28+, Altschul *et al.*, 1990) search among a set of protein coding genes to infer orthogroups and aligns them using MAFFT (v7.123b, Katoh & Standley 2013). These orthogroups may contain paralogs as well as orthologs, and because datasets rich in paralogs can confound phylogenomic analysis, the alignment files produced by OrthoFinder were parsed to recover only those orthogroups that contained single-copy orthologs from each of the 20 species. This resulted in 1,916 total orthogroups with 100% taxon occupancy.

#### Trimming Alignments

For each alignment, regions that contained gap rich sites were removed using –*gappout* option in trimAl v1.4.rev15 (Capella-Gutiérrez *et al.*, 2009). Next, all files containing orthogroups were renamed so the respective headers among these files were identical and individual alignments were concatenated. Concatenation resulted in a single fasta file containing all 1,916 partitions with 1,054,662 sites at 100% taxon occupancy. This initial alignment was further filtered using MARE (v.0.1.2) (Misof *et al.*, 2013), which reduced the character matrix to 247,627 sites. This reduced fasta alignment was converted into a partitioned phylip formatted file. Next, the best-fit substitution models for each partition and a global partitioning scheme were determined with PartitionFinder (v1.1.1) using hcluster clustering algorithm and default parameters (Lanfear *et al.*, 2014).

#### Constructing Phylogeny

Maximum likelihood (ML) analysis was conducted in RaxML v 8.1.20 (Stamatakis 2014) leveraging the partitioning scheme determined by PartitionFinder. The ML tree and 200 bootstrap replicates were performed in a single analysis using the –*f a* option. In addition, we conducted Bayesian Markov Chain Monte Carlo (BMCMC) analysis in MrBayes 3.2.6 (Ronquist *et al.*, 2012). For MrBayes analysis, we specified the mixed amino acid model prior and ran the fully partitioned tree search for 215,000 generations. A consensus tree was then generated after discarding 50% of the run as burnin. The nexus file, including MrBayes block, provides other details of the MrBayes analysis (Methods S1).

#### Detecting genes under positive selection

To identify genes under positive selection in *G. morbida*, we compared *G. morbida* with all nonpathogens from the aforementioned 20 fungi used to estimate the species tree. Among this batch of 10 fungal species, we detected 22,908 protein orthogroups using OrthoFinder that contained paralogs as well as orthologs. Of these, only 9,560 orthogroups were alignable with MAFFT because many groups consisted of only one sequence from a single species (Katoh & Standley, 2013). A total of 3,327 orthogroups, composed of single-copy orthologs, were filtered and corresponding coding DNA sequences for each peptide in these partitions were extracted using custom scripts that can be found <u>online</u>.

The coding DNA sequences were then aligned with MACSE v1.01.b (Ranwez *et al.*, 2011). This Java-based utility accounts for frameshifts and premature stop codons in coding sequences during the alignment process and outputs aligned protein and nucleotide sequences. In order to filter out alignments with frameshifts and internal stop codons, we utilized a program called PAL2NAL v14 (Suyama *et al.*, 2006). This software searches for complementary regions between multiple protein alignments and the corresponding coding DNA sequences, and omits any problematic codons from the output file. This cleaning step reduced the number of 3,327 orthogroups to 2,798 that were used for detecting genes under selective pressures.

We used the branch-site model (BSM) in the CodeML program of package PAML v4.8 for selection analysis (Yang 2007). BSM permits ω (dN/dS) to vary among sites and branches permitting the identification of specific branches and sites subjected to selection. We computed two models in order to calculate and compare the likelihood values: a null model with a fixed ω value of 1 and an alternative model that estimates ω in the foreground branch, which is *G. morbida* in our case. In the effort to reduce false positives, we implemented the Benjamini-Hochberg correction method when comparing likelihood ratios for null and alternative models using a *P*-value threshold of 0.05. We performed similar BLAST searches as mentioned previously to characterize the functions of these proteins and identify proteins with signal peptides and transmembrane domains.

We repeated the above procedures for detecting genes under selection in *Grosmannia clavigera* because this fungal pathogen plays an ecological role similar to *G. morbida.* By performing these analyses, we sought to uncover genes under adaptive evolution in both beetle-vectored tree pathogens.

#### Sequence data and code availability

The raw reads and assembled genomes reported in this paper are available at European Nucleotide Archive under Project Number PRJEB13066. The in silico genereated transcript and protein files are being deposited at Dryad. The code is available at Github (https://github.com/tarunaaggarwal/G.morbida.Comp.Gen)

## Results

### Assembly features

We recently assembled a reference genome for a *G. morbida* strain isolated from *Juglans californica* in Southern California (Schuelke *et al.*, 2016). The reference contained 73 scaffolds with an estimated size of 26.5 Mbp. By using the MAKER annotation pipeline, we predicted 6,273 protein models in this reference in-silico (Cantarel *et al.*, 2008). In this work, we sequenced strains of *G. flava* and *G. putterillii* at approximately 102x and 131x coverage, respectively. The *G. flava* assembly was composed of 1,819 scaffolds totaling 29.47 Mbp in length, and the *G. putterillii* genome contained 320 scaffolds extending 29.99 Mbp. *G. flava* and *G. putterillii* totaled 6,976 and 7,086 protein models, respectively. Both genomes contained 98% of the single-copy orthologs present in more than 90% of the fungal species. Nearly all of the raw reads (97% and 98%) mapped back to *G. flava* and *G. putterillii* genome assemblies, respectively (Table 4). These statistics indicated that our genome assemblies are high quality and complete.

**Table 4.** Length-based statistics for *Geosmithia morbida, Geosmithia flava*, and *Geosmithia putterillii* generated with QUAST v2.3. The average GC content for *G. morbida, G. flava*, and *G. putterillii* equals 54%, 52%, and 55.5% respectively. All genome completeness values were produced with BUSCO v1.1b1. These percentages represent genes that are complete and not duplicated or fragmented.

| Species | Est. genome size (Mbp) | k-mer for ABySS assembly | Scaffold count | Largest scaffold | NG50* | LG50* | Genome completeness | Predicted Proteins |
| --- | --- | --- | --- | --- | --- | --- | --- | --- |
| *G. morbida* | 26.5 | NA<sup>1</sup> | 73 | 2,597,956 | 1,305,468 | 7 | 98 | 6,273 |
| *G. flava* | 29.6 | 91 | 1,819 | 1,534,325 | 460,430 | 22 | 98 | 6,976 |
| *G. putterillii* | 30.0 | 91 | 320 | 2,758,267 | 1,379,352 | 9 | 98 | 7,086 |

<sup>1</sup>Genome assembly for *G. morbida* was constructed using AllPaths-LG (v49414). See Schuelke *et al.*, 2016 for further details. \*NG50 is the scaffold length such that considering scaffolds of equal or longer length produce 50% of the bases of the reference genome. LG50 is the number of scaffolds with length NG50.

An estimated 0.80% of *G. morbida* reference genome sequence represented repeats, whereas 0.63% and 0.64% of the sequences in *G. flava* and *G. putterillii* consisted of repetitive elements. There were 60, 42, and 15 DNA transposons in *G. morbida*, *G. flava*, and *G. putterillii*, respectively. Furthermore, *G. morbida* possessed only 152 retroelements, whereas *G. flava* and *G. putterillii* had 401 and 214 of such elements, correspondingly (Table 5).

**Table 5.** Repetitive elements profile of *Geosmithia* species generated with RepeatMasker v4.0.5.

|  | **Genome size**<br>**(Mbp)** | **GC (%)** | **Bases**<br>**masked**<br>**(%)** | **Retroelements**<br>**(N)** | **DNA**<br>**transposons**<br>**(N)** |
| --- | --- | --- | --- | --- | --- |
| *G. morbida* | 26.5 | 54.30 | 0.81 | 152 | 60 |
| *G. flava* | 29.6 | 51.87 | 0.63 | 401 | 42 |
| *G. putterillii* | 30.0 | 55.47 | 0.64 | 214 | 15 |

Although, the extent to which mobile genetic elements affect genome evolution in *Geosmithia* is unknown, mobile genetic elements may be influential drivers of adaptive evolution in *G. morbida.* They are known to be responsible for genomic rearrangements and expansion, horizontal gene transfer and generation of new genes (Casacuberta & Gonzalez 2013; Stukenbrock & Croll 2014). For example, *Fusarium oxysporum* has a genome nearly 60 Mbp in length and contains 16.83 Mbp of repetitive sequences. *F. oxysporum* also contained four more chromosomes than closely related species, and the added chromosomes were rich in transposons and genes such as putative effectors, necrosis and ethylene-inducing proteins and carbohydrate binding enzymes (Ma *et al.*, 2010). Although *Geosmithia morbida* harbors fewer mobile genetic elements than fungal species such as *F. oxysporum*, it is possible that such elements have contributed to the evolution of pathogenicity in *Geosmithia* via gene expansion and/or horizontal gene transfer. Understanding the role of mobile genetic elements within genus *Geosmithia* may be key in discovering the genetic basis behind the evolution of pathogenicity.

### Identifying putative genes involved in pathogenicity

Approximately 32%, 34%, and 35% of the total proteins in *G. morbida, G. flava* and *G. putterillii* respectively shared significant homology with protein sequences in the database. The number of unknown proteins with hits in the PHI-base database was similar for *G. morbida* (26), *G. flava* (28), and *G. putterillii* (36). The full BLASTp search results against the PHI-base database for *G. morbida, G. flava*, and *G. putterillii* are available in the supporting material (Table S1).

### Identifying species-specific genes

The three *Geosmithia* species and *A. chrysogenum* contained a total of 9065 orthologs and paralogs. Among the set of homologous genes there were 4,655 single copy orthologs. *A. chrysogenum* contained 2338 species-specific genes, of which seven genes were paralogous. *G. morbida* possessed 76 unique genes whereas, *G. putterilli* and *G. flava* had 161 and 146 species-specific genes. The two nonpathogenic *Geosmithia* species did not contain any paralogs, however *G. morbida* had three unique genes present in multiple copies. Based on a functional search against NCBI’s non-redundant database, the three genes encode hydantoinase B/oxoprolinase, aldehyde dehydrogenase, and ABC-2 type transporter.

These findings are significant because all three of these proteins are involved in stress responses that can be induced by the host immune system during the infection process. For example, aldehyde dehydrogenases are part of a large protein family that detoxify aldehydes and alcohols in all organisms including fungal species (Asiimwe *et al.*, 2012). Hydantoinase B/oxoprolinase is involved in the synthesis of glutathione, a compound essential for basic cellular functions but also important in cellular defense against oxidative stress (Pocsi *et al.*, 2004). Glutathione has been shown to chelate damaging metal ions by inhibiting their spread in the cell (Pocsi *et al.*, 2004), and to prevent the accumulation of H_2_O_2_ in *Paxillus involutus* (Ott *et al.*, 2002). Lastly, ATP-binding cassette (ABC) proteins belong to an especially large family of proteins that regulates transport of substances across the cellular membrane. In pathogenic fungi, they are involved in drug resistance and in the production of defense molecules (Krattinger *et al.*, 2009; Wang *et al.*, 2013; Karlsson *et al.*, 2015).

### Identifying putative secreted proteins

A total of 349, 403, and 395 proteins in *G. morbida, G. flava*, and *G. putterillii* contained signal peptides respectively. Of these putative signal peptide-containing proteins in *G. morbida*, 27 (7.7%) encoded proteins with unknown function, whereas *G. flava* and *G. putterillii* contained 29 (7.2%) and 30 (7.6%) unknown proteins, respectively. The difference in percent of unknown proteins with signal peptides was minimal among the three genomes. For each species, proteins containing signal peptides were subjected to a membrane protein topology search using TMHMM v2.0. There were 237, 281, and 283 proteins in *G. morbida, G. flava*, and *G. putterillii* that lacked any transmembrane protein domains. Again, these numbers were not significantly different.

### Profiling carbohydrate active enzymes and peptidases

CAZymes are carbohydrate active enzymes that break down plant structural components, enabling initiation and establishment of infection. We assessed the CAZymatic profile of all species in the order *Hypocreales, Geosmithia* species, and *Grosmannia clavigera* (Figure 1). The glycoside hydrolase (GH) family members dominated all protein models, followed by glycosyltransferase (GT) family. The two most prominent families among all fungal species were GH3 and GH16 (Table S2). GH3 hydrolases are involved in cell wall degradation and overcoming the host immune system, and GH16 enzymes fulfill a wide range of cellular functions including transporting amino acids. The third most representative family was GH18; however *G. morbida* only contained 4 of these enzymes. In contrast, this number for other species ranges from 9 to 31 enzymes. Along with acetylglucosaminidases, family GH18 harbors chitinases that assist in the production of carbon and nitrogen. In terms of other CAZyme families, all fungi except *F. solani* express a similar overall distribution. *Fusarium solani* contains more CAZymes than any other pathogen or non-pathogen. This *Fusarium* species is a generalist necrotrophic pathogen that is believed to possess more CAZymes than biotrophic and hemibiotrophic fungi. This discrepancy may be due to the fact that necrotrophic pathogens require an extensive toolkit to promote host cell death as quickly as possible; whereas biotrophs need to keep the host alive, and dispensing large number of degradation enzymes can be detrimental to that aim (Zhao *et al.*, 2013).

**Figure 1.**
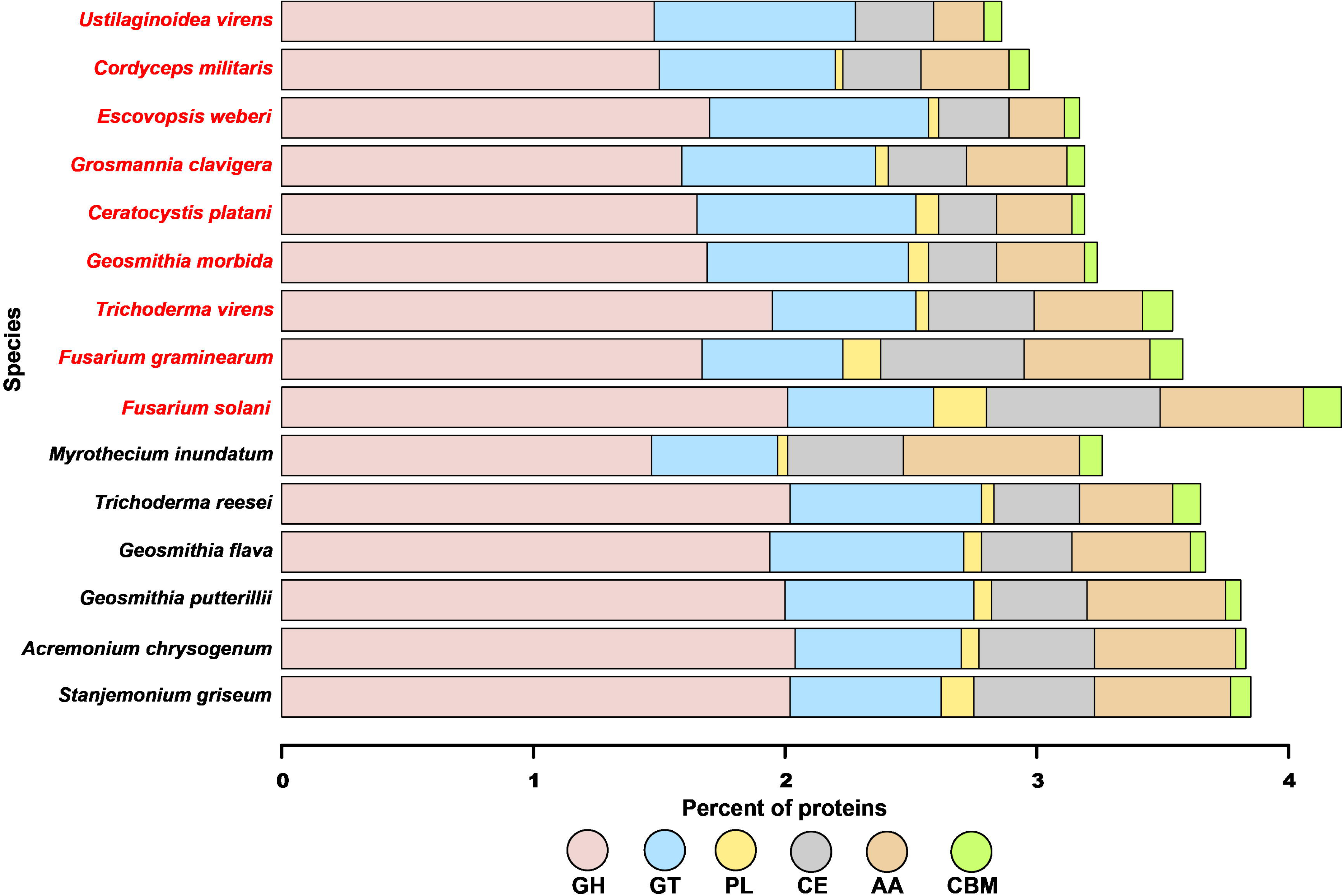
Carbohydrate active enzymes (CAZymes) distribution for *Geosmithia* species, other Hypocreales, and *Ceratocystis platani.* The species in red are pathogens, while the names in black are nonpathogens. CAZymes were identified with HMMer searches of dbCAN peptide models. GH: glycoside hydrolases, GT: glycosyltransferases, PL: polysaccharide lyases, CE: carbohydrate esterases, AA: auxiliary activities enzymes, and CBM: carbohydrate-binding molecules.

In addition to profiling CAZymes, we also performed a BLAST search against the peptidase database—Merops v10.0 (Rawlings *et al.*, 2016) -- for each *Hypocreales, Ceratocystis platani*, and *G. clavigera.* Among the pathogens, *G. morbida* has the third highest percent of predicted proteases after *Cordyceps militaris* (insect pathogen) and *G. clavigera* (Figure 2, Table S3). Moreover, *Geosmithia flava* and *G. putterillii* have the largest percent of peptidases among the nonpathogenic fungi. All three *Geosmithia* species illustrate similar proteolytic profiles and contain no glutamic and mixed peptidases. These results were expected because all three *Geosmithia* species are closely related. Furthermore, given that these species are plant affiliates (except *Cordyceps militaris)*, the ability to degrade lignin and cellulose is an important life history trait that is conserved throughout fungal pathogens, but perhaps did not give rise to pathogenicity in *G. morbida*.

**Figure 2.**
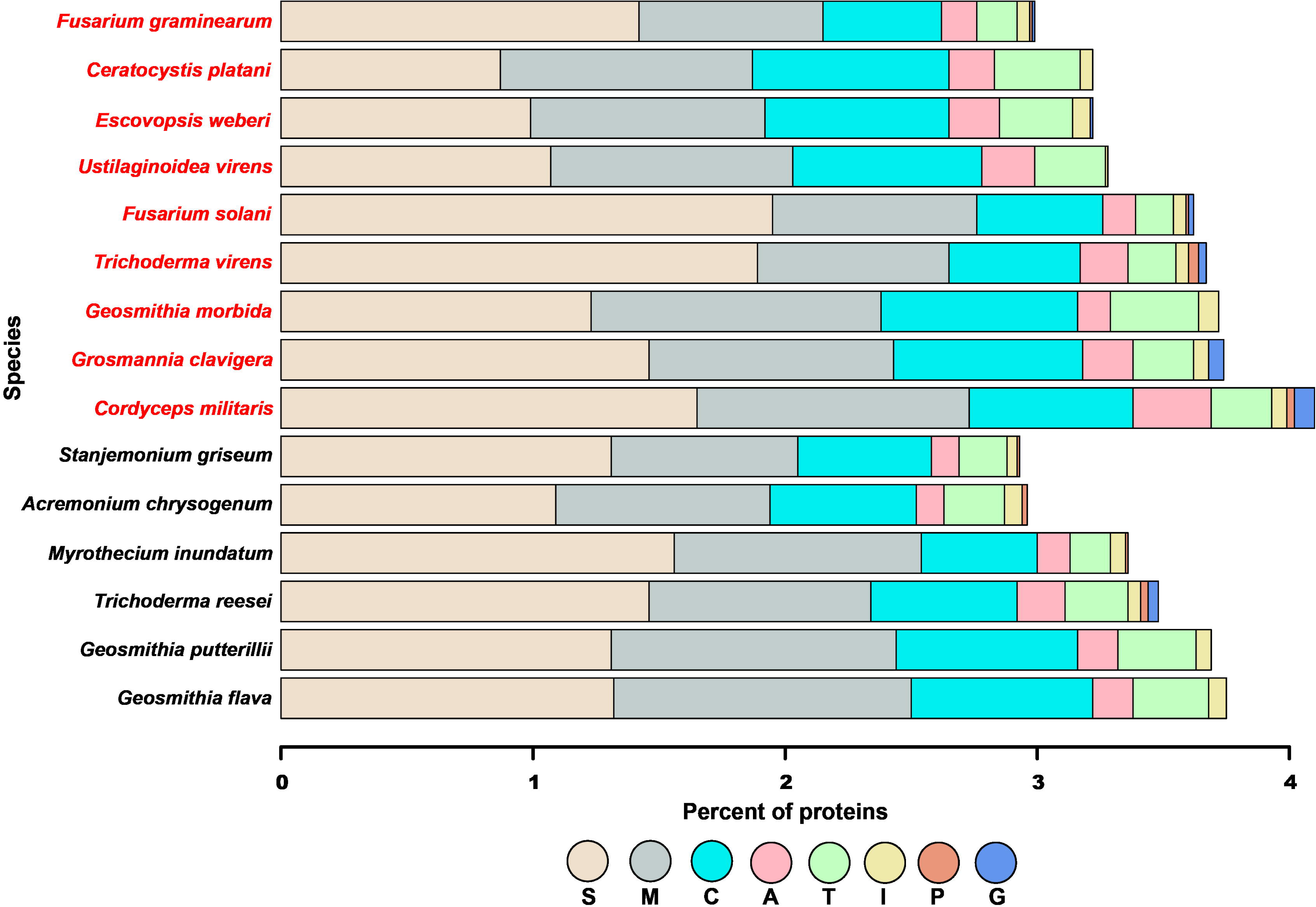
Proteolytic enzymes distribution for *Geosmithia* species, other Hypocreales, and *Ceratocystis platani.* The species in red are pathogens, while the names in black are nonpathogens. Proteases were identified using BLASTp searches against the MEROPs database v10. S: serine, M: metallo, C: cysteine, A: aspartic, T:threonine, I: inhibitors, P: mixed, G: glutamic.

### Inferring phylogeny

Even though *Geosmithia* was first established as a genus in 1979, it has only recently been described in depth. One of the main objectives in this study was to uncover the phylogenetic relationship between *Geosmithia* species and other fungal pathogens using coding DNA sequence data. In order to determine the broader evolutionary history of *Geosmithia* species, we constructed maximum likelihood (ML) and Bayesian Markov Chain Monte Carlo (BMCMC) phylogenies using 1,916 single-copy orthologs from *G. morbida, G. putterillii, G. flava*, and 17 additional fungal taxa (Table 3). Our final dataset consisted of 11 pathogens and 9 non-pathogens. After trimming and filtering, our 1,916 orthogroups contained approximately 1e10^6^ amino acid sites in total. The topologies of trees generated under ML and BMCMC were identical, and all nodes in all analyses received bootstrap support of 100% (ML) and posterior probabilities of 1.0 (BMCMC). The analyses resulted in a single, identical tree topology (Figure 3) that was supported by previous research (Fitzpatrick *et al.*, 2006, Wang *et al.*, 2009). *Geosmithia* species form a monophyletic clade with two nonpathogenic fungi—*Acremonium chrysogenum* and *Stanjemonium griseum*—indicating that the common ancestor shared among these species was not a pathogen.

**Figure 3.**
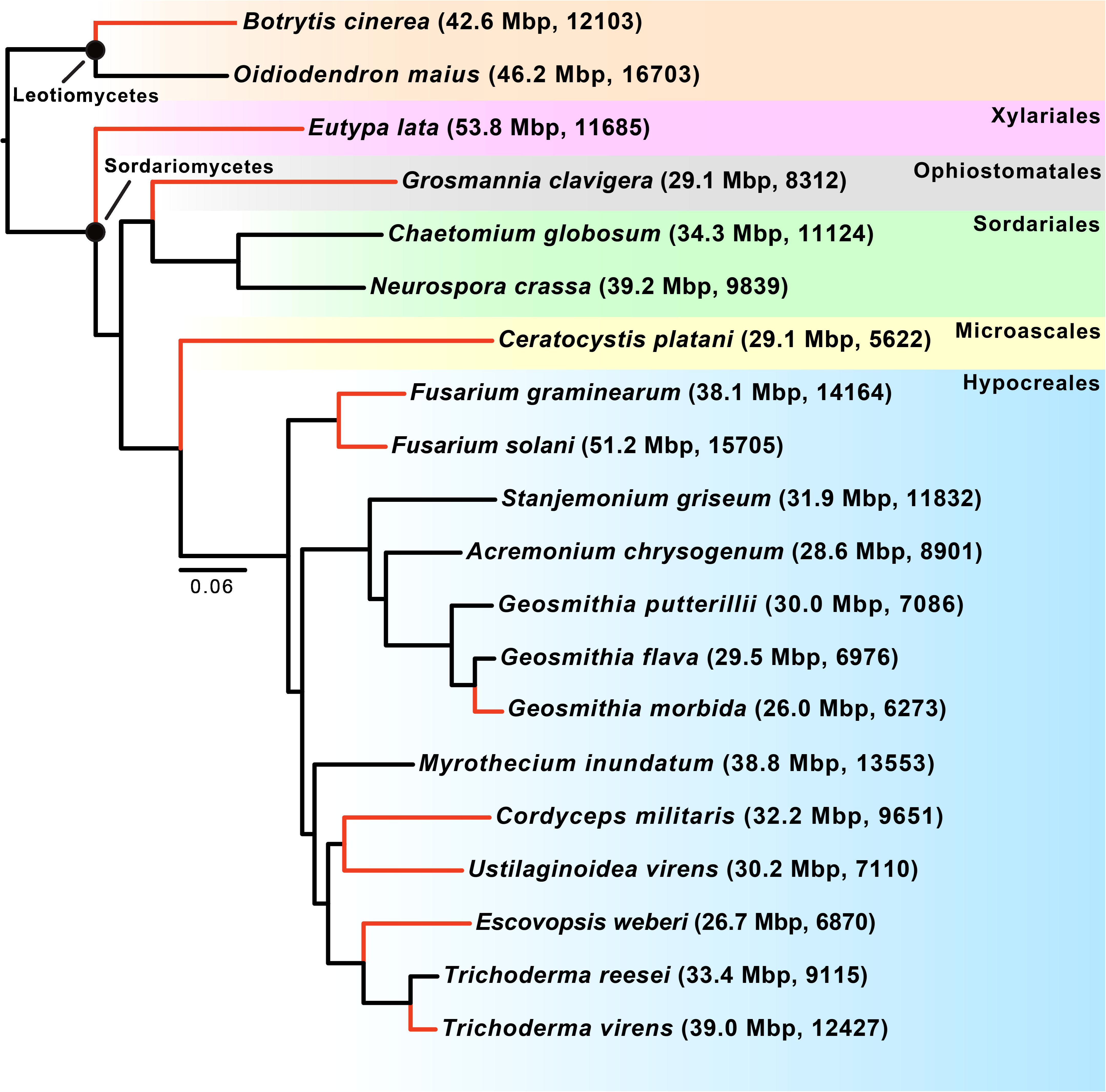
The Bayesian Markov Chain Monte Carlo (BMCMC) phylogeny was estimated using the mixed amino acid model in MrBayes (Ronquist *et al.*, 2012) on a dataset containing 89,999 positions. This topology is identical to partitioned analyses conducted in RAxML (Stamatakis 2014). All nodes in BMCMC and ML analyses receive maximum support. The black circles symbolize classes. The color-shaded boxes at the right of the figure denote the orders within each class. The first and second numbers in parentheses represent the genome sizes in Mbp and the number of predict protein models, respectively. Black and red branches correspond to non-pathogens and pathogens, respectively, which span multiple orders.

### Genes under positive selection

In order to understand the molecular basis of pathogenicity in *G. morbida*, we sought to detect genes under positive selection. For this, we first searched for all single-copy orthologs shared among the 9 non-pathogens and *G. morbida* using OrthoFinder (v0.3.0). We extracted the corresponding coding sequences for each protein in the 3,327 orthogroups containing 1:1 orthologs using a custom python script. These orthogroups were aligned using MACSE v1.01b and cleaned with PAL2NAL v14. After alignment and cleaning of orthogroups there were 2,798 multiple sequence alignments that were used for selection analysis.

To identify coding sequences and sites under selection, we leveraged the branch-site model in PAML’s codeml program (4.8). *Geosmithia morbida* was selected as the foreground branch. Our results showed 38 genes to be under positive selection using an adjusted *P*-value < 0.05. Next, we performed a functional search for each protein by blasting the peptide sequences against the NCBI non-redundant and pfam databases. We determined that several were involved in catabolic activity, gene regulation, and cellular transport (Table 6, Table S4).

**Table 6.**
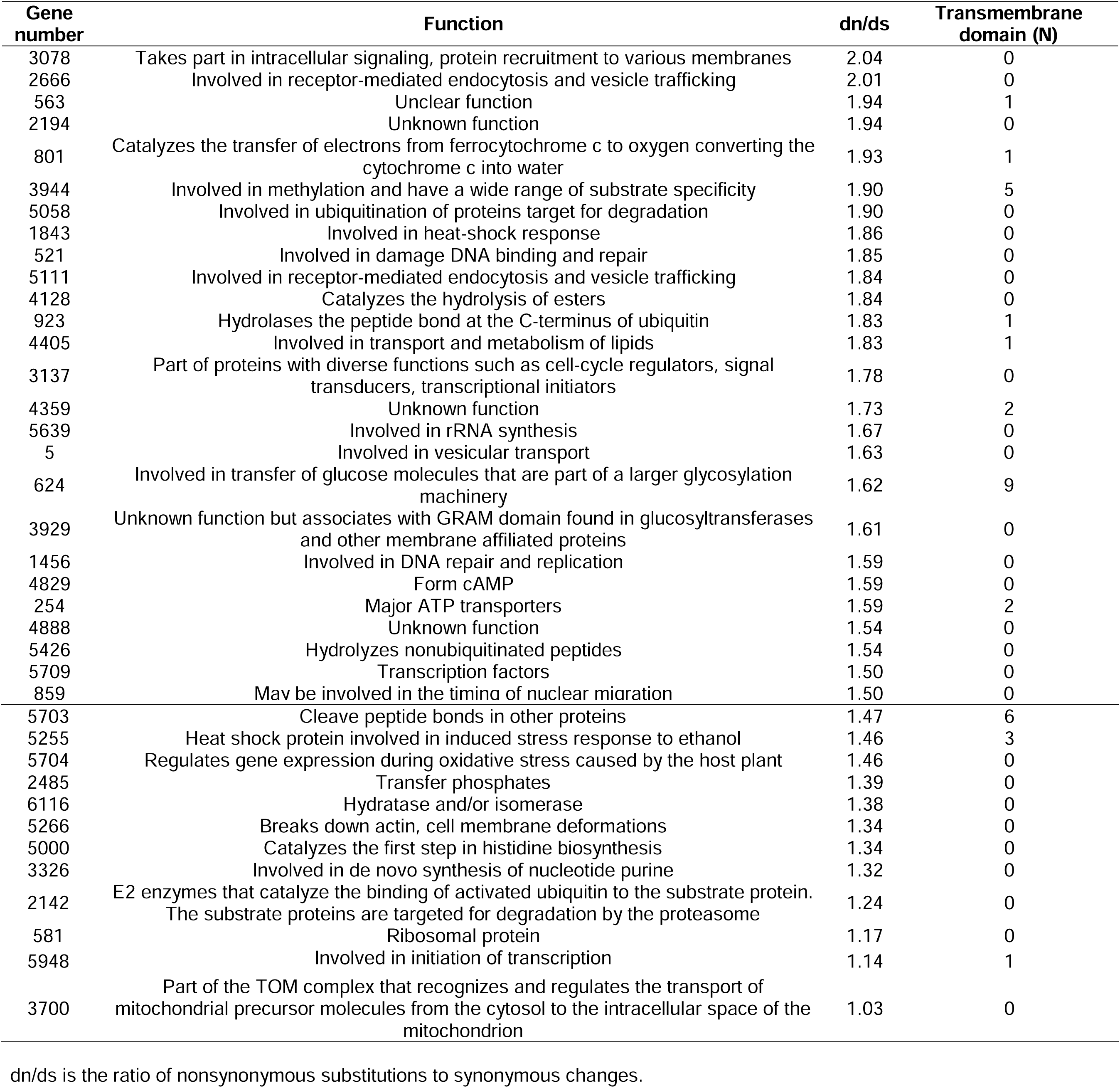
Functional analyses of genes under positive selection in *Geosmithia morbida* detected by the branch-site model in PAML 4.8. The gene number corresponds to the sequence ID in the *G. morbida* protein file available at DRYAD.

| Gene number | Function | dn/ds | Transmembrane domain (N) |
| --- | --- | --- | --- |
| 3078 | Takes part in intracellular signaling, protein recruitment to various membranes | 2.04 | 0 |
| 2666 | Involved in receptor-mediated endocytosis and vesicle trafficking | 2.01 | 0 |
| 563 | Unclear function | 1.94 | 1 |
| 2194 | Unknown function | 1.94 | 0 |
| 801 | Catalyzes the transfer of electrons from ferrocytochrome c to oxygen converting the cytochrome c into water | 1.93 | 1 |
| 3944 | Involved in methylation and have a wide range of substrate specificity | 1.90 | 5 |
| 5058 | Involved in ubiquitination of proteins target for degradation | 1.90 | 0 |
| 1843 | Involved in heat-shock response | 1.86 | 0 |
| 521 | Involved in damage DNA binding and repair | 1.85 | 0 |
| 5111 | Involved in receptor-mediated endocytosis and vesicle trafficking | 1.84 | 0 |
| 4128 | Catalyzes the hydrolysis of esters | 1.84 | 0 |
| 923 | Hydrolases the peptide bond at the C-terminus of ubiquitin | 1.83 | 1 |
| 4405 | Involved in transport and metabolism of lipids | 1.83 | 1 |
| 3137 | Part of proteins with diverse functions such as cell-cycle regulators, signal transducers, transcriptional initiators | 1.78 | 0 |
| 4359 | Unknown function | 1.73 | 2 |
| 5639 | Involved in rRNA synthesis | 1.67 | 0 |
| 5 | Involved in vesicular transport | 1.63 | 0 |
| 624 | Involved in transfer of glucose molecules that are part of a larger glycosylation machinery | 1.62 | 9 |
| 3929 | Unknown function but associates with GRAM domain found in glucosyltransferases and other membrane affiliated proteins | 1.61 | 0 |
| 1456 | Involved in DNA repair and replication | 1.59 | 0 |
| 4829 | Form cAMP | 1.59 | 0 |
| 254 | Major ATP transporters | 1.59 | 2 |
| 4888 | Unknown function | 1.54 | 0 |
| 5426 | Hydrolyzes nonubiquitinated peptides | 1.54 | 0 |
| 5709 | Transcription factors | 1.50 | 0 |
| 859 | May be involved in the timing of nuclear migration | 1.50 | 0 |
| 5703 | Cleave peptide bonds in other proteins | 1.47 | 6 |
| 5255 | Heat shock protein involved in induced stress response to ethanol | 1.46 | 3 |
| 5704 | Regulates gene expression during oxidative stress caused by the host plant | 1.46 | 0 |
| 2485 | Transfer phosphates | 1.39 | 0 |
| 6116 | Hydratase and/or isomerase | 1.38 | 0 |
| 5266 | Breaks down actin, cell membrane deformations | 1.34 | 0 |
| 5000 | Catalyzes the first step in histidine biosynthesis | 1.34 | 0 |
| 3326 | Involved in de novo synthesis of nucleotide purine | 1.32 | 0 |
| 2142 | E2 enzymes that catalyze the binding of activated ubiquitin to the substrate protein.<br>The substrate proteins are targeted for degradation by the proteasome | 1.24 | 0 |
| 581 | Ribosomal protein | 1.17 | 0 |
| 5948 | Involved in initiation of transcription | 1.14 | 1 |
| 3700 | Part of the TOM complex that recognizes and regulates the transport of<br>mitochondrial precursor molecules from the cytosol to the intracellular space of the<br>mitochondrion | 1.03 | 0 |

dn/ds is the ratio of nonsynonymous substitutions to synonymous changes.

For instance, a cullin3-like protein was predicted to be under positive selection. Cullin3-like proteins belong to a group of structurally similar molecules involved in protein degradation, such as the Skp-Cullin-F-box (SCF) ubiquitin ligase complex, was predicted to be under positive selection (Cardozo & Pagano 2004; Pintard *et al.*, 2004). Furthermore, a ubiquitin-conjugating enzyme (E2) that interacts with cullin3 to prepare substrate for degradation, also had a dn/ds > 1, indicating that both genes are under positive selection within *G. morbida.* Although little is known regarding the precise functional abilities of these complexes, it is possible these proteins are involved in pathogenicity of *G. morbida.* Previous studies have also implicated ubiquitin ligase complexes in infection and disease development (Duyvesteijn *et al.*, 2005; Han *et al.*, 2007).

Our analysis also revealed a regulatory protein homologous to basic leucine zipper (bZIP) transcription factors was under selection. The bZIP proteins are similar to AP-1 transcription factors and monitor several developmental and physiological processes including oxidative stress responses in eukaryotes (Corrêa *et al.*, 2008). Fungal pathogens such as the rice blast fungus *Magnaporthe oryzae* express AP1-like transcription factor called MoAP1 that contains bZIP domain. MoAP1 is highly active during infection and is translocated from the cytoplasm to the nucleus in response to oxidative stress induced by H_2_O_2_ (Guo *et al.*, 2011). MoAP1 regulates enzymes such as laccase and glutamate decarboxylase that are involved in lignin breakdown and metabolism of γ-aminobutyric acid, respectively (Solomon & Oliver 2002; Baldrian 2005; Janusz *et al.*, 2013). Some of the other positively selected genes include ABC transporter, proteases, proteins involved in apoptosis, and proteins related to DNA replication and repair. As previously mentioned, ABC transporters are important mediators that aid in protection against plant defenses as well as natural toxic compounds (Krattinger *et al.*, 2009; Wang *et al.*, 2013; Karlsson *et al.*, 2015; Lo Presti *et al.*, 2015). Apoptosis or programmed cell death helps establish resistance during host-microbe interactions, helps organisms cope with oxidative environments, and may even be essential for infection (Veneault-Fourrey *et al.*, 2006; Kabbage *et al.*, 2013). In fungal species, proteins involved in DNA replication and repair are essential for the formation and penetration of appressorial structures into the host cell (Son *et al.*, 2016). Only five of the 38 genes with evidence of selection encoded proteins with unknown functions. These positively selected genes might be involved in the evolution and adaptation of *G. morbida*.

### Transmembrane protein and effector genes

Transmembrane proteins are important mediators between a host and its pathogens during microbial invasion. Fungal pathogens either penetrate a surface or enter the host through a wound or opening such as stomata in order to gain access to the nutrients in the plant (Chisholm *et al.*, 2006). Once the infiltration process is completed, pathogens are exposed to host plasma membrane receptors that detect pathogen-associated molecular patterns (PAMP) and induce PAMP-triggered immunity (PTI) to prevent further proliferation of the microbe. Transmembrane proteins expressed by a fungal pathogen are crucial during PTI because they are responsible for suppressing PTI directly or by secreting effector molecules, which contain signal peptides necessary for proper targeting and transport (Chisholm *et al.*, 2006; Boller & He 2009). Our analysis of the 38 proteins under positive selection showed that 11 of these possess at least one or more transmembrane domains. Although nearly 30% of the positively selected genes identified were membrane bound, a similar proportion of non-selected genes in *G. morbida* were membrane associated, so this result is not strong evidence that interactions with the host surface are drivers of evolution within *G. morbida.* Among proteins under selection we found no protein that contained a signal peptide, indicating none of these proteins are secretory.

### Genes under adaptive evolution in beetle-vectored fungal pathogens

In addition to detecting genes under selective pressures in *G. morbida*, we performed the same selection analysis for *Grosmannia clavigera* to identify overlapping proteins that may help explain adaptations leading to the ecological role these two beetle-vectored fungi play. We found that *G. clavigera* possessed 42 positively selected genes that shared protein domains with only two of the 38 genes predicted to be under selection in *G. morbida.* The two overlapping motifs are methyltransferase and protein kinase domains. Our KEGG analysis exhibited no common pathways between *G. morbida* and *G. clavigera.* These findings emphasize that evolutionary forces act differently on divergent populations. All fungal pathogens face dissimilar environmental challenges and associate with different hosts both spatially and temporally. Even closely related organisms can be highly distinct molecularly. For instance, the fungi responsible for the Dutch elm disease—*Ophiostoma ulmi* and *O. novo-ulmi*—differ in their genetic composition and virulence despite their strong evolutionary relationship (Brasier 2001; Khoshraftar *et al.*, 2013; Comeau *et al.*, 2015).

## Discussion

This study aims to provide insight into the evolution of pathogenicity within *Geosmithia morbida*, a beetle vectored pathogen that is the causal agent of Thousand Cankers Disease in *Julgans* species. Here, we present *de novo* genome assemblies of two nonpathogenic *Geosmithia* species, *G. flava* and *G.* putterillii, and employ comparative genomics approach to uncover the molecular factors contributing to pathogenicity in *G. morbida.*

*G. flava* and *G. putterillii* have estimated genome sizes of 29.6 Mbp and 30.0 Mbp, correspondingly. These assemblies are larger than the genome of *G. morbida*, which measures 26.5 Mbp in length. In contrast to other species in the phylogeny (Figure 3), fungi associated with trees either as pathogens or saprophytes *(Geosmithia* species, *G. clavigera*, and *C. platani)* had reduced genomes and gene content. We predict this genome and gene content reduction is a result of evolving specialized lifestyles to occupy a specialized niche. For instance, all three *Geosmithia* species and *G. clavigera* are vectored into their respective hosts via bark beetles, which may result in strong selection on the genetic variability of the fungi because they must adapt to their vectors and hosts simultaneously. Moreover, possessing genes that are not essential for this specialized lifestyle may impose a fitness disadvantage on the pathogen, as they may represent potential targets for host resistance genes. A recent study characterizing the genome of mycoparasite *Escovopsis weberi* showed that specialized pathogens tend to have smaller genomes and predicted protein sets because they lack genes that are not required beyond their restricted niche when compared to closely related generalists (de Man *et al.*, 2016). Our results agree with this finding because *G. morbida* has a more specialized beetle vector *(P. juglandis)* and plant host range *(Juglans* species) in comparison to *G putterilli* and *G. flava* which can be found on a variety of trees species including both gymnosperms and angiosperms (Kolařík *et al.*, 2004; Kolařík & Jankowiak 2013), and can be vectored by multiple beetle species (Kolařík *et al.*, 2008.) This represents a significant contrast to previous reports that have documented the importance of genome expansion with the evolution of pathogenicity (Adhikari *et al.*, 2014; Raffaele & Kamoun 2012). Furthermore, our results are supported by prior findings which showed that gene loss and gain can lead to a more specialized lifestyle in bacterial and eukaryotic lineages (Ochman & Moran 2001, Lawrence 2005). Another example is a study by Nagendran *et al.*, 2009 that showed *Amanita bisporigera*, which is an obligate ectomycorrhizal symbiont, lacks many plant cell-wall-degrading enzymes suggesting that these genes may no longer be required for *A. bisporigera’s* specialized lifestyle.

Genome reduction is an important evolutionary mechanism that propels divergence of species and more often than not enables adaptation to specific environments. Although genome reduction is more frequent in prokaryotes, it is not uncommon among eukaryotes including fungal species (Ochman and Moran 2001, Nagendran *et al.*, 2009, Spanu *et al.*, 2010).

Although one might expect the pathogen *G. morbida* to possess more carbohydrate binding enzymes and peptidases than its non-pathogenic relatives, our results indicated that all three species had similar enzymatic profiles (Figures 1 and 2). Despite these similarities, our PAML analysis identified 38 genes under positive selection in *G. morbida* when compared to other nonpathogens within the order Hypocreales including non-pathogenic *Geosmithia* species. These genes encode for proteins that have been implicated in pathogenicity in other fungal pathogens such as *Magnaporthe oryzae.* Additionally, we found peptides with protein kinase and methyltransferase domains that are under positive selection in both *G. morbida* and *G. clavigera.* Proteins kinases were previously shown to be under strong positive selection in *G. clavigera* (Alamouti *et al.*, 2014). This result was especially important given the key contributions that protein kinases make in initiating signal transduction pathways during pathogen host interactions. Our study identified a small set of genes potentially involved in the evolution of pathogenicity in the genus *Geosmithia.* Functional experiments and analyses of the expression levels of these genes during infection as compared to gene expression of a non-pathogen would shed light on the mechanisms influencing pathogen evolution and genome reduction.

## Acknowledgments

The use of trade names is for the information and convenience of the reader and does not imply official endorsement or approval by the United States Department of Agriculture or the Forest Service of any product to the exclusion of others that may be suitable. Partial funding was provided by the New Hampshire Agricultural Experiment Station. Special thanks to Dr. Miroslav Kolarik for providing isolates of *Geosmithia flava* and *Geosmithia putterilli.* We are also grateful to Dr. Joseph Spatafora and his team for giving us permission to utilize sequence data for *Stanjemonium griseum* and *Myrothecium inundatum.* Lastly, we thank the 1000 Fungal Genomes Project for being a valuable source of genetic data.

## Author contribution

TAS conceived, designed and performed the experiments and wrote the manuscript. AW assembled and annotated the genomes. KB conceived and designed the study, wrote and reviewed the manuscript. KB also conceived funding. DCP designed and implemented the phylogenetic methods in this study and reviewed the manuscript. KW conveived funding and designed the experiments. KW also wrote and reviewed the manuscript. MDM conceived and designed the study, developed analyses pipelines and edited the manuscript.

## Support Information

The following Supporting Information is available for this article:

**Table S1** Complete BLASTp results against the Phibase4 database for *Geosmithia morbida, Geosmithia flava* and *Geosmithia putterillii.*

**Table S2** CAZymes hits data for 20 fungal species used in this study.

**Table S3** Merops hits data for 20 fungal species used in this study

**Table S4** Complete BLASTp results against NCBI’s nr database for 38 genes found to be under selection in *Geosmithia morbida.*

**Methods S1** A modified nexus file illustrating Mr. Bayes block used for constructing phylogeny.

